# scMerge: Integration of multiple single-cell transcriptomics datasets leveraging stable expression and pseudo-replication

**DOI:** 10.1101/393280

**Authors:** Yingxin Lin, Shila Ghazanfar, Kevin Wang, Johann A. Gagnon-Bartsch, Kitty K. Lo, Xianbin Su, Ze-Guang Han, John T. Ormerod, Terence P. Speed, Pengyi Yang, Jean Yee Hwa Yang

## Abstract

Concerted examination of multiple collections of single cell RNA-Seq (scRNA-Seq) data promises further biological insights that cannot be uncovered with individual datasets. However, such integrative analyses are challenging and require sophisticated methodologies. To enable effective interrogation of multiple scRNA-Seq datasets, we have developed a novel algorithm, named scMerge, that removes unwanted variation by combining stably expressed genes and utilizing pseudo-replicates across datasets. Analysis of large collections of publicly available datasets demonstrates that scMerge performs well in multiple scenarios and enhances biological discovery, including inferring cell developmental trajectories.

## Introduction

Single-cell transcriptome profiling by next generation sequencing (scRNA-Seq) has enabled unprecedented resolution in studying cell identity, heterogeneity, and differentiation trajectories in various biological systems^1^. Comprehensive characterisation of large collections of scRNA-Seq datasets can provide a more holistic understanding of the underlying biological processes. However, the integration of multiple scRNA-Seq datasets remains a challenge due to prevailing technical effects associated with experiments performed across multiple conditions, experiments, and organisms. Here, we have developed scMerge, an algorithm for integrative biological analysis using multiple scRNA-Seq datasets. scMerge corrects for specific batch effects within an experiment as well as removing dataset-specific effects across collections of datasets.

While normalization methods such as SCnorm^2^, scran^3^, mnnCorrect^4^, and ComBat^5^ can be applied for combining multiple scRNA-Seq datasets, they are either not specifically designed for adjusting batch effects, or are primarily designed in the context of removing batch effects within a single experiment. Alternatively, data integration methods such as Seurat^6^, fastMNN^4^, and ZINB-WaVE^7^ generate dimension reduced datasets where individual genes cannot be examined for downstream analysis such as differential expression based marker identification or pseudotime trajectory estimation. Moreover, with the exception of mnnCorrect^4^, these existing methods make an underlying assumption that the batches or datasets to be integrated contain the same or similar proportions of particular cell types. This assumption can lead to incorrectly normalized data, especially when particular batches or datasets have markedly different relative proportions of cell types, e.g. integrating two datasets where one dataset contains fluorescence activated cell sorted cells and the other does not. mnnCorrect aims to address this by estimating a set of ‘mutual nearest neighbors’, a mapping of distinct individual cells between batches or datasets, but can be unstable due to the selection of these individual pairs of cells, as opposed to selection of pairs of cell clusters.

## Methods

To enable effective integration of multiple scRNA-Seq datasets, we have developed a novel algorithm called scMerge. The scMerge algorithm consists of three key components (Fig. 1a):

i. the identification of stably expressed genes (scSEGs) via a Gamma-Gaussian mixture model^8^ for use as ‘negative controls’ for estimating unwanted factors;
ii. the construction of pseudo-replicates to estimate the effects of unwanted factors; and
iii. the adjustment of the datasets with unwanted variation using a fastRUVIII model.

**Figure 1.**
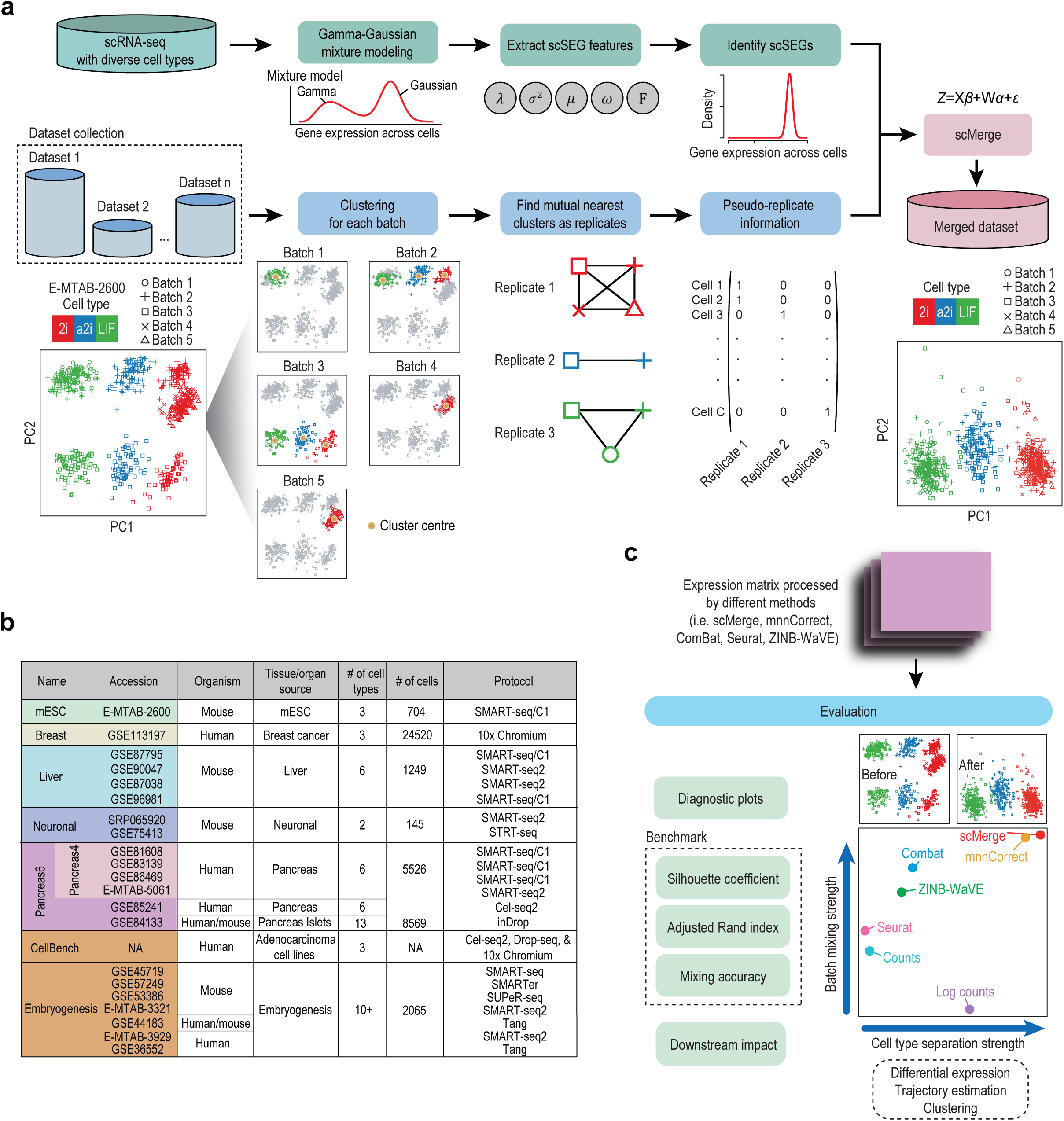
Schematic illustration of the scMerge algorithm. (a)First, stably expressed genes (scSEGs) are identified using a reference dataset with diverse cell types, to be used as negative control genes. Second, for a given data collection with multiple datasets, clustering is performed per dataset, mutual nearest clusters are identified across datasets. Selected cells from these clusters are then identified as pseudo-replicates, to be treated as replicates in the factor analysis step. Factor analysis is performed with the negative control genes and pseudo-replicates, resulting in a single merged dataset. (b)Summary of 14 datasets comprising six data collections used in this study. (c)Summary of evaluation strategies for merged datasets using diagnostic plots, indices comparing to known cell type labels and further downstream impacts.

Details of these components can be found in the Online Methods. scMerge takes gene expression matrices from a collection of datasets and a list of negative control genes whose expressions are expected to be constant across these datasets. The final output is a single normalized and batch corrected gene expression matrix with all input matrices merged ready for further downstream analysis. scMerge is available online as an R package at github.com/SydneyBioX/scMerge.

## Evaluation

To assess the performance of scMerge for integrating multiple scRNA-Seq datasets, we collated fourteen publicly available scRNA-Seq datasets into six distinct data collections (Fig. 1b, Supplementary Table 1). Each data collection varies across key characteristics, including number of datasets, sequencing platforms, species, as well as cell type compositions (Fig. 1b). We compared scMerge to other approaches, including scran^3^, mnnCorrect^4^, ComBat^5^, Seurat^6^ and ZINB-WaVE^7^. The performance of each method was evaluated using multiple criteria including visual inspection of diagnostic plots, Silhouette coefficients, adjusted Rand indices (ARI), as well as downstream biological impact (Fig. 1c and Online Methods).

Using a mixture modelling approach^8^, our algorithm defines stably expressed genes (1076 for human and 826 for mouse) that are characterized by low variability and wide range of expression (Fig. 2a). Expression of scSEGs show minimal association with cell types and developmental stages compared to previously identified housekeeping genes from bulk transcriptome data (bHK)^9,10^ or random subsets of genes (n = 1076; Fig. 2b). We found that, in cases where batch labels are unknown and thus pseudo-replicates cannot be identified, using scSEGs as negative control genes results in better integration of data (Fig. 2c, Supplementary Fig. 10), compared to bHK genes. Consistent with this, we found that using scSEGs as negative controls also results in better integration of data (F1-scores) than using ERCC spike-ins controls (Supplementary Fig. 11), potentially due to the exogenous nature of ERCC probes. Conceptually, the choice of scSEGs will have a greater effect when integrating heterogeneous datasets with large differences between cells and a high proportion of highly variable genes, and as a result, appropriate selection of negative control genes has a large influence on the normalization results.

**Figure 2.**
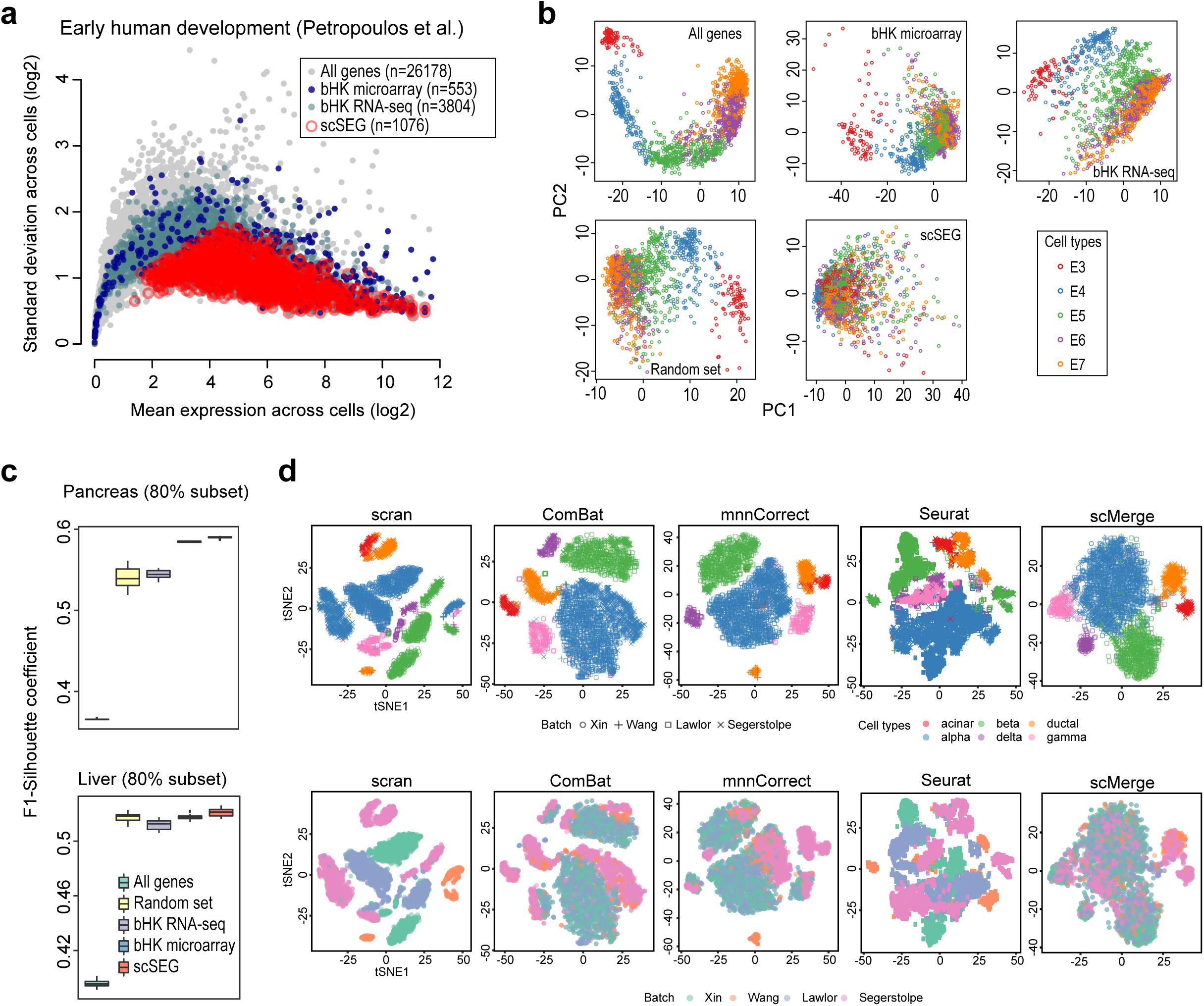
Characterising single cell stably expressed genes (scSEGs) (a)Scatter plot showing mean expression (x-axis) and standard deviation (y-axis) on log scale of each gene (grey circles) across profiled single cells. Open red circles represent stably expressed genes (scSEGs) derived for human in this study whereas dark and light blue solid circles represent housekeeping genes defined previously using bulk microarray (bHK microarray)^9^ and RNA-Seq (bHK RNA-Seq)^10^ data. (b)A panel of principal component analysis (PCA) plots based on all genes or different subset of genes including, bHK microarray, bHK RNA-Seq data, scSEGs as well as a random selection of genes, for the Petropoulos et al^16^ data. (c)A 2 by 1 panel of boxplots comparing the effect of different types of negative controls for the Pancreas4 data collection (top panel) and Liver data collection (bottom panel). The y-axis represents the F1-score of Silhouette coefficients between cell type mixing and (1 - datasets mixing), where higher values are desired. Stratified sampling is performed to randomly subset 80% of cells from the datasets, repeated 10 times to examine stability. (d)A 2 x 5 panel of tSNE plots of the Pancreas4 data collection using the output from scran, ComBat, mnnCorrect, Seurat, and scMerge (using scSEGs as negative controls). The top row of is color coded by cell types and the second row is color coded by the distinct Pancreas datasets.

We compared scMerge with four popular and recent batch correction methods: ComBat^5^, mnnCorrect^4^, Seurat^6^ and ZINB-WaVE^7^ (Online Methods and Fig. 1c) using our six scRNA-seq data collections that cover different tissues, species and protocols (Fig. 1b). We found that scMerge effectively removed batch and dataset specific effects across a wide range of biological systems, including a collection of human pancreatic scRNA-Seq datasets (Fig. 1b). Visual inspection of tSNE plots (Fig. 2d), and similarly for PCA plots (Supplementary Fig. 1), shows that unlike other methods, scMerge clearly separates acinar and ductal cells. Additionally, in scMerge processed datasets, cell type information explained a higher percentage of ‘wanted’ variation than ‘unwanted’ variation^11^ (Supplementary Fig. 13).

In general, we found that scMerge performed favourably in terms of maintaining strong biological signal and reducing unwanted variation such as batch or data specific noise (Fig. 3a, Supplementary Fig. 1-8, and Supplementary Fig. 14). Our evaluation metrics (Online Methods) capture the trade-off between these two broad objectives. scMerge manages the trade-off between separating cell types and merging batches well (Fig. 3a) across multiple data collections in comparison with other methods. Summarizing these two quantities into a single F1-score (Online Methods), we found that scMerge maintains this better performance, despite choice of Silhouette coefficient or ARI as the comparison metric (Supplementary Fig. 9, Supplementary Table 2).

**Figure 3.**
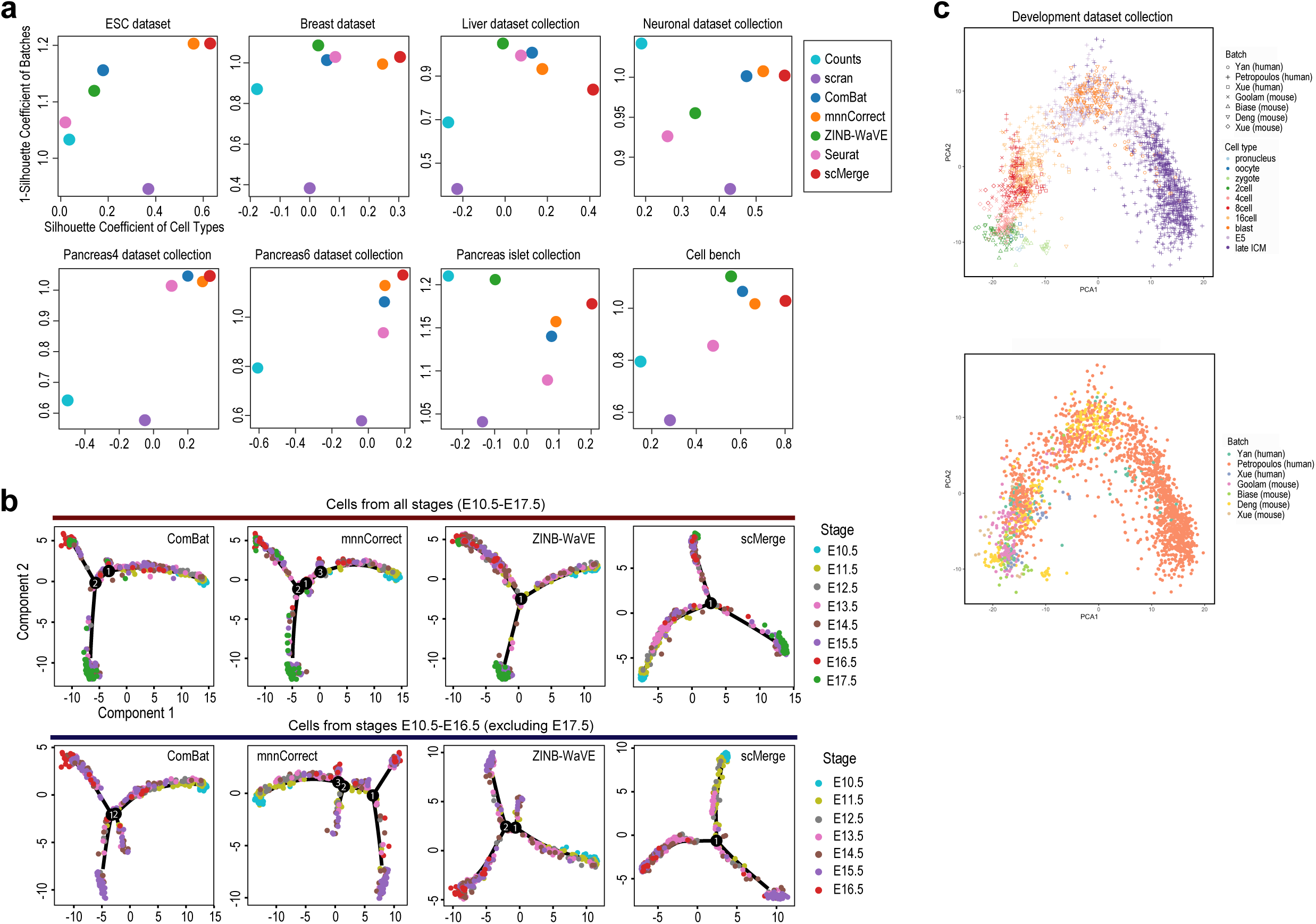
Comparison results. (a)A 2 by 4 panel of scatter plots of Silhouette coefficients for no normalization (Counts), scran, ComBat, mnnCorrect, ZINB-WaVE, Seurat, and scMerge (using scSEGs as negative controls). The x-axes denote the Silhouette coefficient of cell types and y-axes denote the 1 – Silhouette coefficient of batch effects, where desirable outcomes are in the top-right hand corner. (b)A 2 by 4 panel of pseudotime trajectories demonstrating the stability of scMerge. The first row displays the trajectories from Monocle 2 using hepatoblasts, hepatocytes, and cholangiocytes from all time points in the Liver data collection, and the second row displays the trajectories from Monocle 2 with the Liver data collection with time point E17.5 removed. (c)A 2 by 1 panel of PCA plots of the Embryogenesis data collection following scMerge (using scSEGs as negative controls). The top panel is color coded by cell types and the bottom panel is color coded by the seven different ESC datasets.

To illustrate the capability of scMerge to enable further downstream analyses, we studied the integrated expression matrices of the Liver data collection: four liver scRNA-seq datasets taken from different experimental settings (Fig. 1b). To examine stability in the face of incomplete data, we reconstructed the cell trajectories, using Monocle 2^12^, of hepatoblasts, hepatocytes, and cholangiocytes for both the full Liver data collection and for a subset of the original Liver data collection, where cells corresponding to the E17.5 time point of GSE90047^13^ were removed. We found that the trajectory associated with scMerge is most consistent with the full Liver data collection (Fig. 3b) and agrees with current literature^14^ (Supplementary Fig. 15), while other methods tended to generate extraneous branches with the subset of the Liver data collection.

Finally, we illustrated the potential of scMerge in facilitating fine-grained annotation of cell types during early human and mouse development by integrating the Embryogenesis data collection: seven datasets that profiled human^15^-^17^ and mouse^18^-^20^ embryogenesis at various stages ranging from zygotes to late blastocysts (Fig. 1b). By matching the zygote, 2-cell, 4-cell, 8-cell, and 16-cell stages across these datasets with semi-supervised scMerge (Online Methods), we observed clear time-course separation of cells from zygotes to late blastocysts (Fig. 3c), as well as clear overlap of many cells from human and mouse blastocysts, inner cell mass (ICM) regions, and human embryonic day 5 (E5). Because these three categories each contain multiple cell types, similar in their development and lineage specification stages, scMerge processed data therefore allows different cell types in these categories to be matched across species based on their relationship in the dimension reduced space.

In conclusion, scRNA-Seq technology permits a cell-type specific characterization of gene expression, enriching our understanding of the underlying biological processes. Examination of effectively integrated collections of scRNA-Seq datasets promises further biological insights that may not be possible from analyzing each individual dataset. Here we have shown that scMerge allows investigators to achieve this goal and has a significant downstream impact on the inference of cell trajectories. Integration of large collections of embryogenesis datasets across human and mouse illustrate the utility of scMerge in annotating cell types in development across different species.

## Methods

Methods, including statements of data availability and any associated accession codes and references, are available in the online version of the paper. The scMerge R package is available at github.com/SydneyBioX/scMerge.

## Author contributions

JYHY conceived the study with input from SG, KW, and PY. YL led the method development and data analysis with input from SG, KW, JTO, JAG, TPS, PY and JYHY. JYHY, PY, YL and SG interpreted the results with input from KW, JAG, TPS, XS and ZGH. KKL curated the data with input from YL and JYHY. YL and KW implemented the R package with input from JAG and JTO. JYHY, PY, YL, SG and KW wrote the manuscript with input from KKL and TPS. All authors read and approved the final version of the manuscript.

### Acknowledgements

The authors thank all their colleagues, particularly at The University of Sydney, School of Mathematics and Statistics, for their support and intellectual engagement. The following sources of funding for each author, and for the manuscript preparation, are gratefully acknowledged: Australian Research Council Discovery Project grant (DP170100654) to JYHY, JTO, PY; Discovery Early Career Researcher Award (DE170100759) to PY; Australia NHMRC Career Developmental Fellowship (APP1111338) to JYHY and the Judith and David Coffey Life Lab at the Charles Perkins Centre, The University of Sydney to SG. Australian Postgraduate Award to KW. Research Training Program Tuition Fee Offset and Stipend Scholarship to YL. NHMRC Program Grant (1054618) to TPS. J.G. was supported by the National Science Foundation under grant no. DMS-1646108. SJTU-USYD Translate Medicine Fund-Systems Biomedicine AF6260003. The funding source had no role in the study design; in the collection, analysis, and interpretation of data, in the writing of the manuscript, and in the decision to submit the manuscript for publication.

## References

1. Jaitin, D. A. et al. Massively Parallel Single-Cell RNA-Seq for Marker-Free Decomposition of Tissues into Cell Types. Science (80-.). 343, 776–779 (2014).

2. Bacher, R. et al. SCnorm: robust normalization of single-cell RNA-seq data. Nat. Methods 14, 584–586 (2017).

3. Lun, A. T. L., McCarthy, D. J. & Marioni, J. C. A step-by-step workflow for low-level analysis of single-cell RNA-seq data with Bioconductor. F1000Research 5, 2122 (2016).

4. Haghverdi, L., Lun, A. T. L., Morgan, M. D. & Marioni, J. C. Batch effects in single-cell RNA-sequencing data are corrected by matching mutual nearest neighbors. Nat. Biotechnol. 36, 421–427 (2018).

5. Johnson, W. E., Li, C. & Rabinovic, A. Adjusting batch effects in microarray expression data using empirical Bayes methods. Biostatistics 8, 118–27 (2007).

6. Butler, A., Hoffman, P., Smibert, P., Papalexi, E. & Satija, R. Integrating single-cell transcriptomic data across different conditions, technologies, and species. Nat. Biotechnol. 36, 411–420 (2018).

7. Risso, D., Perraudeau, F., Gribkova, S., Dudoit, S. & Vert, J.-P. A general and flexible method for signal extraction from single-cell RNA-seq data. Nat. Commun. 9, 284 (2018).

8. Ghazanfar, S., Bisogni, A. J., Ormerod, J. T., Lin, D. M. & Yang, J. Y. H. Integrated single cell data analysis reveals cell specific networks and novel coactivation markers. BMC Syst. Biol. 10, 11–24 (2016).

9. Eisenberg, E. & Levanon, E. Y. Human housekeeping genes are compact. Trends Genet. 19, 362–365 (2003).

10. Eisenberg, E. & Levanon, E. Y. Human housekeeping genes, revisited. Trends Genet. 29, 569–574 (2013).

11. McCarthy, D. J., Campbell, K. R., Lun, A. T. L. & Wills, Q. F. Scater: pre-processing, quality control, normalization and visualization of single-cell RNA-seq data in R. Bioinformatics btw777 (2017). doi:10.1093/bioinformatics/btw777

12. Qiu, X. et al. Reversed graph embedding resolves complex single-cell trajectories. Nat. Methods 14, 979–982 (2017).

13. Yang, L. et al. A single-cell transcriptomic analysis reveals precise pathways and regulatory mechanisms underlying hepatoblast differentiation. Hepatology 66, 1387–1401 (2017).

14. Müsch, A. From a common progenitor to distinct liver epithelial phenotypes. Curr. Opin. Cell Biol. 54, 18–23 (2018).

15. Yan, L. et al. Single-cell RNA-Seq profiling of human preimplantation embryos and embryonic stem cells. Nat. Struct. Mol. Biol. 20, 1131–9 (2013).

16. Petropoulos, S. et al. Single-Cell RNA-Seq Reveals Lineage and X Chromosome Dynamics in Human Preimplantation Embryos. Cell 165, 1012–1026 (2016).

17. Xue, Z. et al. Genetic programs in human and mouse early embryos revealed by single-cell RNA sequencing. Nature 500, 593–7 (2013).

18. Goolam, M. et al. Heterogeneity in Oct4 and Sox2 Targets Biases Cell Fate in 4-Cell Mouse Embryos. Cell 165, 61–74 (2016).

19. Biase, F. H., Cao, X. & Zhong, S. Cell fate inclination within 2-cell and 4-cell mouse embryos revealed by single-cell RNA sequencing. Genome Res. 24, 1787–96 (2014).

20. Deng, Q., Ramsköld, D., Reinius, B. & Sandberg, R. Single-cell RNA-seq reveals dynamic, random monoallelic gene expression in mammalian cells. Science 343, 193–6 (2014).

